# MUSE-XAE: MUtational Signature Extraction with eXplainable AutoEncoder enhances tumour type classification

**DOI:** 10.1101/2023.10.23.562664

**Authors:** Corrado Pancotti, Cesare Rollo, Giovanni Birolo, Piero Fariselli, Tiziana Sanavia

## Abstract

Mutational signatures are a critical component in deciphering the genetic alterations that underlie cancer development and have become a valuable resource for understanding the genomic changes that occur during tumorigenesis. In this paper, we present MUSE-XAE, a novel method for mutational signature extraction from cancer genomes using an explainable Auto-Encoder. Our approach employs a hybrid architecture consisting of a nonlinear encoder that can capture nonlinear interactions and a linear decoder, ensuring the interpretability of the active signatures in cancer genomes. We evaluated and compared MUSE-XAE with other available tools on synthetic and experimental cancer datasets and demonstrated that it achieves very accurate extraction capabilities while enhancing tumour-type classification. Our findings indicate that the use of Auto-Encoders is feasible and effective. This approach could facilitate further research in this area, with neural network-based models playing a critical role in advancing our understanding of cancer genomics

## 1. Introduction

Mutational signatures are patterns of somatic mutations which reflect the underlying biological processes that drive cancer development [1, 2, 3]. Mutational signatures have become a valuable resource for deciphering the genetic alterations underpinning cancer and for developing targeted therapies [4, 5, 6]. In recent years, many methods have been developed for mutational signature extraction, the majority of which are based on matrix factorization techniques, such as non-negative matrix factorization (NMF) [7, 8, 9, 10, 11, 12, 13] and its probabilistic versions [14, 15, 16]. These approaches have generated significant success across various types of cancer, identifying more than 60 distinct signatures associated with specific mutational processes [17, 18]. Although these techniques have proven to be highly effective for the extraction of mutational signatures, some studies have highlighted possible issues that may arise during extraction that deserve attention and further investigation [19, 20]. Many of the mutational signatures found in the COSMIC catalog are currently without known etiology. Some are merely statistically linked to a specific process, whereas others lack any statistical association. Additionally, the high degree of similarity between certain signatures may suggest the existence of non-biological, overfitted signals [21, 22, 23]. Future extraction techniques should pay heed to these observations, implementing constraints in the decomposition to reduce the detection of overly similar profiles. Furthermore, the intrinsically linear nature of NMF could be a limitation as it might not capture potential nonlinear dependencies within the genome that might contribute to cancer development. To overcome these challenges, we present MUSE-XAE, MUtational Signatures Extraction with eXplainable AutoEncoder. This model includes a nonlinear encoder and a linear decoder with non-negative constraint and a minimum volume regularization [24], adept at capturing potential nonlinear dependencies while preserving signature interpretability. AutoEncoders have been successfully implemented across various domains, including genomics, to glean compact and informative data representations. In particular, autoencoders employing a hybrid architecture with a non linear encoder and a linear decoder has been applied in the context of single-cell RNA-seq and trascriptomic data [25, 26], achieving great success due to their explainability while preserving powerful performance capabilities. However, from the best of our knowledge, this is the first application of such architecture in the context of mutational signatures analysis.

To fill this gap, in this paper we present MUSE-XAE and demonstrate its effectiveness on various cancer datasets by comparing it to existing state-of-the-art approaches in both synthetic scenarios and real-world applications, by analyzing the PCAWG dataset. Our method emerges as one of the best-performing approaches in signature extraction and significantly enhances tumor type classification.

## 2. Materials and methods

### MUSE-XAE architecture

An AutoEncoder is a type of neural network able to learn a lower dimensional representation of the data. Given an input space *X* it consists of an encoder network *f*, represented by one or more layers, that maps the input data to a lower-dimensional latent space *Z*, and a decoder network *g* that reconstructs the input space from the latent representation. The goal of an AutoEncoder is to minimize the reconstruction error 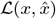 between the original input *x* and the reconstructed output 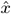. In particular, the general equations that define an AutoEncoder are:

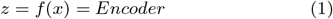

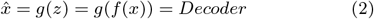

Typically, both functions *f* and *g* are nonlinear activation functions.

MUSE-XAE implements a hybrid architecture with a nonlinear encoder for learning a latent representation *z* of cancer samples and a linear decoder, with non negative constraint and minimum volume regularization to reconstruct the original input, such as 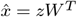. More specifically MUSE-XAE encoder *f* is composed by three hidden layers with batch normalization and a softplus activation function. The decoder *g* is constituted by a weight matrix *W* with non negativity constraint and linear activation function that ensures the interpretability and a minimum volume regularization that helps the model to find a more disentangled representation.

In addition MUSE-XAE exploits a non negative Poisson likelihood function to better take into consideration the count nature of the input data and an early stopping criteria to avoid over fitting. Considering all the contributions the total loss function 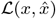 can be written as:

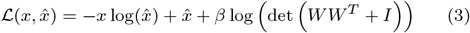

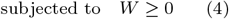

Referring to mutational signatures terminology, the latent representation *z* represents cancer genomes exposures while the decoder weight matrix *W* represents the mutational signatures. MUSE-XAE architecture is sketched in Fig.1.

**Figure 1:**
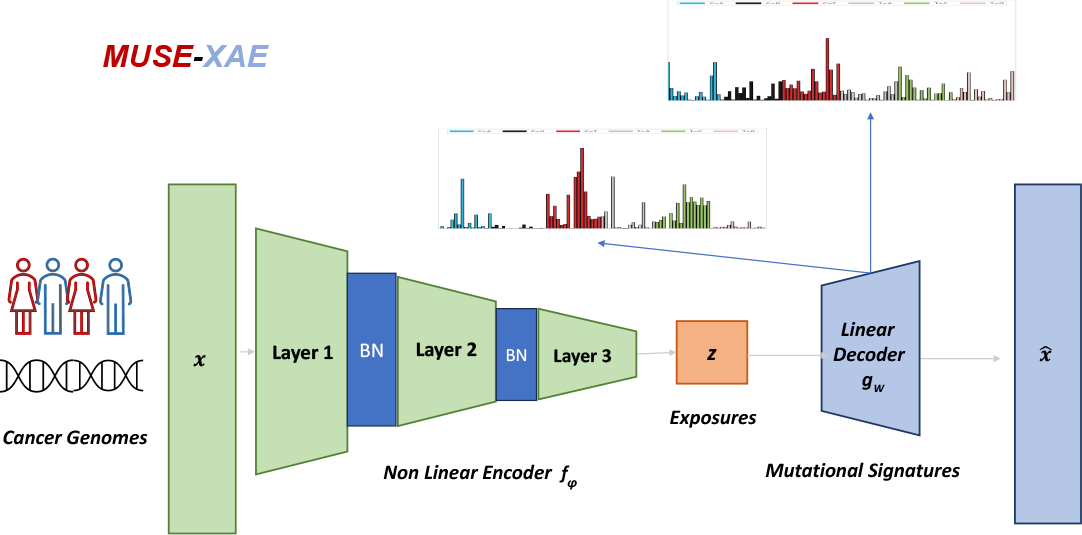
MUSE-XAE schematic architecture. MUSE-XAE features a nonlinear encoder, made up of from three layers that leverage a softplus activation function with batch normalization. The decoder is designed to be linear to enhance interpretability.

### De-Novo Extraction

Training a neural network requires a substantial amount of data to exploit its capacity and fit the parameters effectively. We used a data augmentation strategy to overcome this challenge for the De-Novo extraction of mutational signatures. Specifically, given a tumour catalogue matrix *C* ∈ *R*^*m×*96^, where *m* is the number of samples and 96 is the number of mutational channels, for each cancer genome with *N* total number of mutations, we determined the relative mutation frequency *p* for each of the 96 mutational classes. Then, we generated new data points by bootstrapping cancer genomes *t* times, using a multinomial distribution *M* (*N, p*), obtaining the augmented count matrix *C*_*aug*_.

The approach of bootstrapping cancer genomes, sampling from a multinomial distribution, is already used by other tools to ensure the stability of a consensus signature [1, 7, 17]. We repeat this process *t* times to increase the dataset size.

Then to select the optimal number of active signatures *K*, we used a revised version of the NMFk approach, originally described by [27] and also adopted by SigProfilerExtractor.

Therefore for each number of candidate signatures *k*, MUSE-XAE was trained *n* times. Subsequently a custom KMeans clustering with matching, using cosine similarity distance, was performed on the set of the decoder weights matrices *{W*_1*k*_…*W*_*nk*_*}*, to find a consensus signatures matrix *S*_*k*_. The custom K-Means clustering uses the Jonker-Volgenant algorithm [28] to solve the linear assignment problem in order to find *k* clusters of equal size *n*.

Finally, we considered only the solutions with mean and minimum silhouette scores above a fixed threshold. Among these solutions, the optimal is considered the one with the lowest reconstruction error. MUSE-XAE De-Novo Extraction procedure is summarized in Algorithm 1.

#### Algorithm 1: De-Novo Extraction procedure

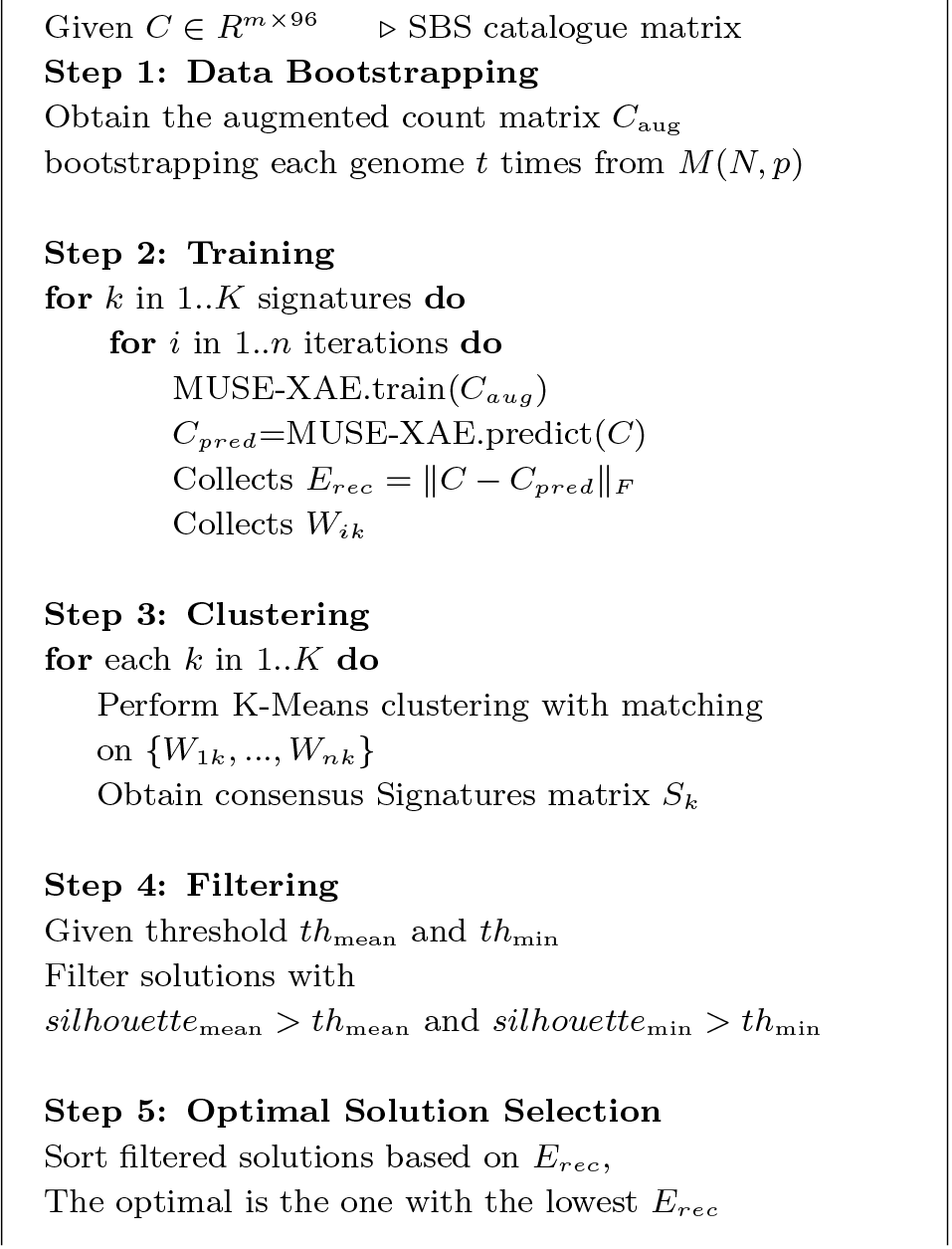

All the parameters mentioned above can be specified by the user, including the size factor *t* for data augmentation, the number of repetition *n* for each candidate signature *k* and the thresholds for the mean and minimum silhouette scores, *th*_mean_ and *th*_min_.

A version with suggested default parameters based on experiments is provided at https://github.com/compbiomedunito/MUSE-XAE.

### Signatures Assignment

Once the profiles of active signatures within a set of cancerous genomes have been identified, it is necessary to understand which signatures are causing mutations within a genome and in what quantity, i.e., we need to assign the contribution of each extracted signature to each genome. To accomplish this, we utilize a slightly modified version of MUSE-XAE used for the signatures extraction. Specifically, once computed the consensus matrix *S*_*k*_, we normalize it, obtaining 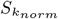and use this matrix to initialize the weights of the decoder and then freeze them, meaning the decoder is no longer trainable. We only allow the weights of the encoder to be trained. To obtain a sparser representation and to avoid over-assignment of mutational signatures, we use an L1 penalty for both the weights of the last layer of the encoder and for the output of the encoder after a ReLU activation, and train the network until convergence. Our new latent representation *z* represents the exposure of the signatures within the genomes, i.e., the number of mutations of a certain mutational class that a signature causes within a genome. We summarized the signature assignment procedure in the Algorithm 2.

#### Algorithm 2: Signature Assignment Procedure

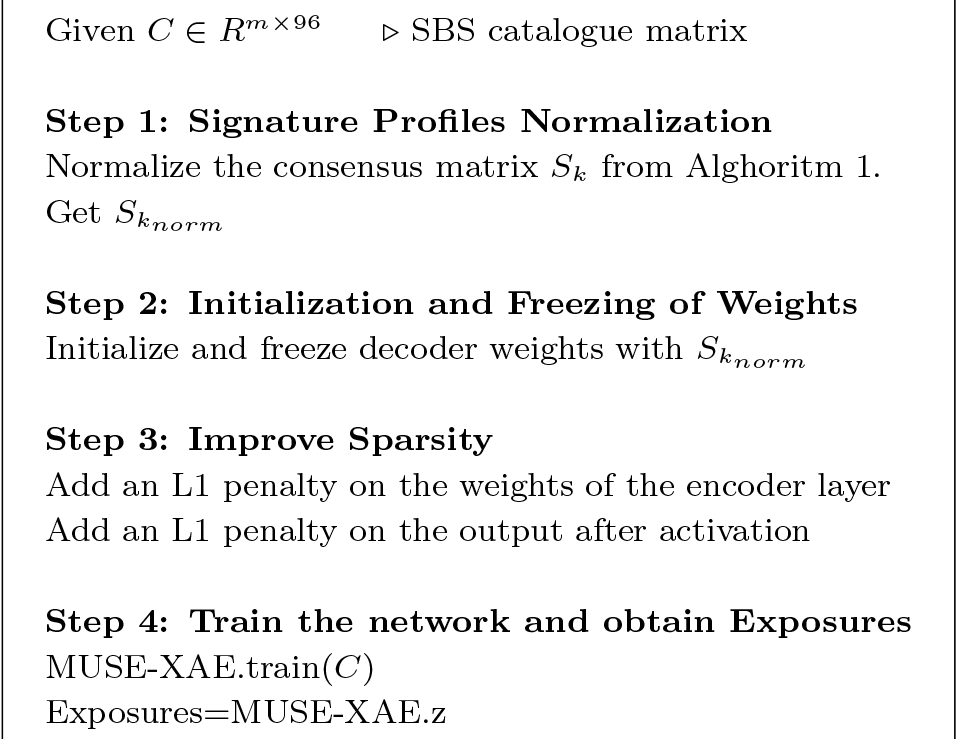

### De-Novo Extraction Scenarios

To evaluate the performance of MUSE-XAE in the context of mutational signature extraction, we utilized 5 publicly available (ftp://alexandrovlab-ftp.ucsd.edu/pub/publications/Islam_et_al_SigProfilerExtractor/) realistic synthetic scenarios also used in the thorough benchmarking conducted in the SigProfilerExtractor paper [7] for currently available mutational signature tools. Specifically, we used the following scenarios:

- **Scenario 1:** 1,000 synthetic samples, modeling a subset of the pancreatic adenocarcinoma PCAWG dataset. The 11 ground-truth signatures are based on COSMIC.
- **Scenario 2:** 1,000 synthetic tumors created from flat, relatively featureless mutational signatures. It includes a mix of 500 synthetic renal cell carcinomas (high prevalence and mutation load from SBS5 and SBS40 signatures) and 500 synthetic ovarian adenocarcinomas (high prevalence and mutation load from SBS3), with a total of 11 COSMIC-based signatures.
- **Scenario 3:** 1,000 synthetic tumors created from signatures with overlapping and potentially interfering profiles, mostly SBS2, SBS7a, and SBS7b. The mutational load distributions were drawn from bladder transitional cell carcinoma (SBS2) and skin melanoma (SBS7a, SBS7b), with 11 COSMIC-based signatures.
- **Scenario 4:** 1,000 synthetic tumors emulating a mix of 500 synthetic renal cell carcinomas (high prevalence and mutation load from SBS5 and SBS40 signatures) and 500 synthetic ovarian adenocarcinomas (high prevalence and mutation load from SBS3). In this scenario, only 3 COSMIC-based signatures (SBS3, SBS5, SBS40) are present.
- **Scenario 5:** 2,700 synthetic samples with mutational spectra matching the ones observed in PCAWG, including 300 spectra from each of 9 different cancer types: bladder transitional cell carcinoma, esophageal adenocarcinoma, breast adenocarcinoma, lung squamous cell carcinoma, renal cell carcinoma, ovarian adenocarcinoma, osteosarcoma, cervical adenocarcinoma, and stomach adenocarcinoma. The ground-truth signatures are 21 signatures based on COSMIC.

Specifically, we extracted mutational signatures from each of these scenarios using MUSE-XAE and applied the same performance metrics as in [7]. We used the Hungarian algorithm to match the predicted and the known signatures based on the cosine-similarity scores. Given that the signatures in each scenario are known, an extracted signature is considered correctly identified, or a True Positive (TP), if the cosine similarity between the extracted and real signature is *>*= *threshold*. If the profile of a signature is missing, it is considered a False Negative (FN), and if the cosine similarity is *< threshold* it is considered a False Positive (FP).

We calculated metrics for Precision, Recall, and the F1 Score for each scenario from the corresponding confusion matrices. Specifically, we computed each metric by varying the cosine similarity threshold from 0.8 to 1. The Precision, Recall, and F1-score are defined as follows:

- Precision is the proportion of True Positives out of the sum of True Positives and False Positives:

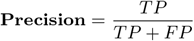

- Recall (also known as Sensitivity) is the proportion of True Positives out of the sum of True Positives and False Negatives:

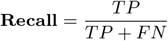

- F1 Score is the harmonic mean of Precision and Recall, which can be expressed mathematically as:

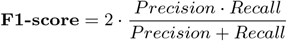

For each measure of accuracy (Precision, Sensitivity and F1), we evaluated the score as a function of the cosine similarity threshold. We then computed their Area Under the Curve (AUC). Since the cosine similarity threshold ranges from 0.8 to 1, the maximum AUC value is 0.2 (*AUC*_*max*_), we normalized the AUC score by dividing by as follows:

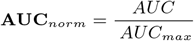

This normalization allowed us to provide a more comparable and interpretable measure of performance across different methods and scenarios.

### PCAWG analysis

To better understand the capacity of MUSE-XAE, we also applied the method to the real Pan Cancer Analysis of Whole Genome (PCAWG) cohort containing 2,780 tumors of 37 cancer types. Specifically, we conducted the following analyses:

1. De Novo extraction of mutational signatures and comparison of profiles with ones of SigProfilerExtractor and with ones known from COSMIC.
2. Analysis of how discriminative the signatures and consequently the exposures are, performing a multi-class classification of cancer types. Specifically, we used the exposures as new features that we fed into a Random Forest to classify both the 25 primary tissue types and the 37 cancer subtypes.

We evaluated the metrics of balanced accuracy, MCC (Matthews Correlation Coefficient), and Kohen Kappa score in a 5-fold cross-validation setting. The computed metrics are defined as follows:

- **Balanced Accuracy**: It is the average recall obtained on each class:

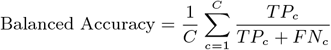

where *T P*_*c*_ and *FN*_*c*_ are the number of true positives and false negatives for the *c*-th class, and *C* is the total number of classes.

- **Multiclass Matthews Correlation Coefficient (MCC)**: It is defined as:

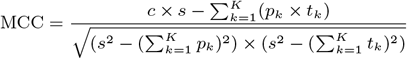

where 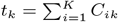 is the number of times class *k* truly occurred, 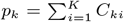 is the number of times class *k* was predicted, 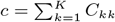 is the total number of samples correctly predicted, and 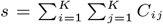 is the total number of samples.

- **Cohen’s Kappa**: This score measures the agreement between two raters who each classify N items into C mutually exclusive categories. The formula for the Kappa score is:

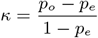

where *p*_*o*_ is the relative observed agreement among raters, and *p*_*e*_ is the hypothetical probability of chance agreement.

## 3. Results

### Data augmentation

In our initial analysis, we investigated the influence of data augmentation on the extraction of mutational signatures across each of the five synthetic scenarios. Specifically, we conducted De-Novo extraction with MUSE-XAE for every dataset, adjusting the data augmentation level from 1 to 100 times the original size. We repeated the extraction five times at each augmentation level to evaluate stability and accuracy.

As depicted in Fig. 2, for all five datasets, there is a trend that an increase in data augmentation not only enhances run-to-run stability, but also improves the count of extracted signatures and their alignment with the real number of profiles.

**Figure 2:**
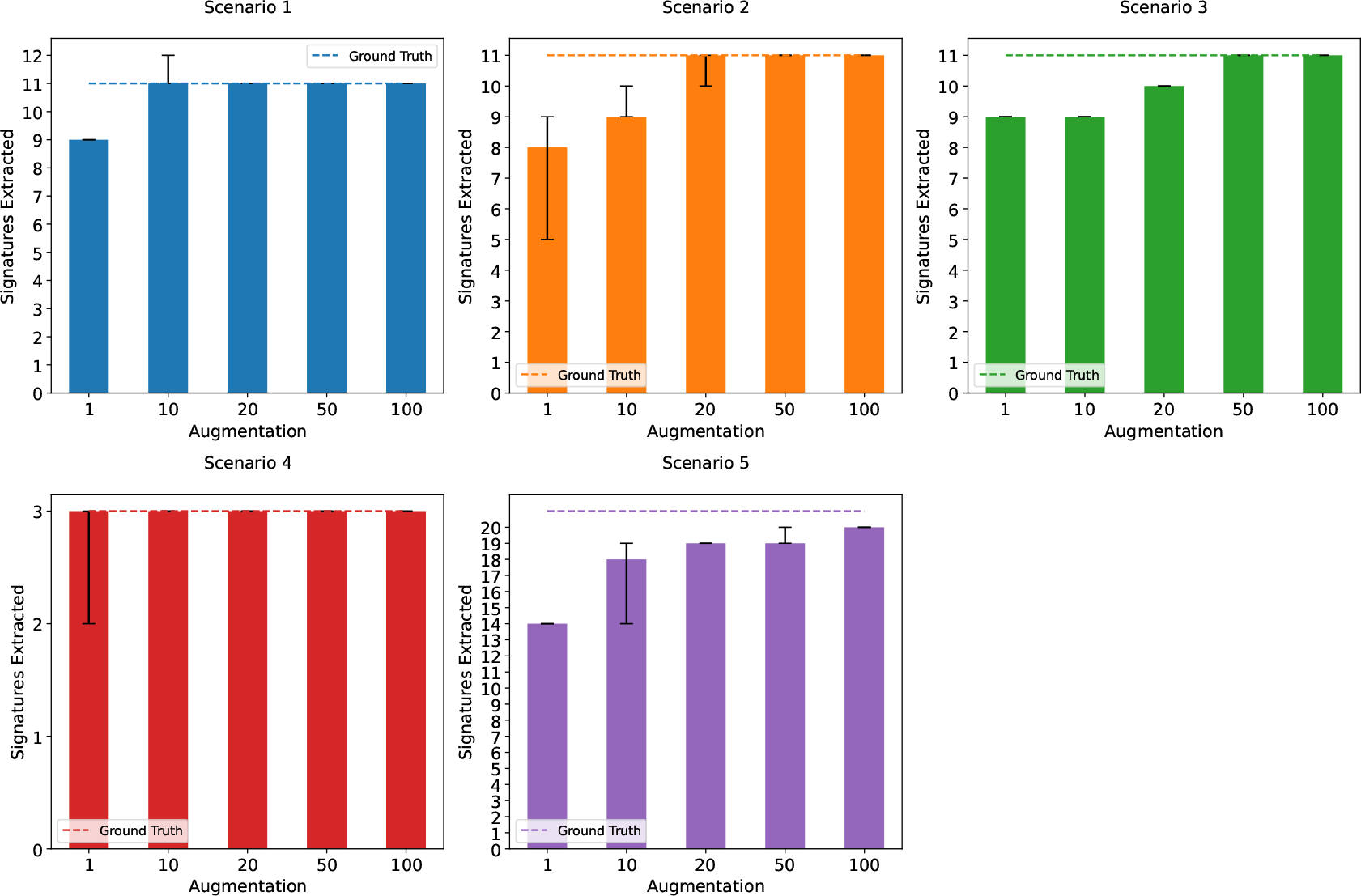
Sensitivity analysis for data augmentation for each of the five synthetic scenario. Each bar represents the mean number of extracted signatures over 5 repetitions. The dashed line represents the ground truth, while the black error bar represents the minimum and maximum.

While data augmentation by itself does not definitively ascertain the accuracy of real signature profile extraction, it nonetheless offers valuable insights. It gives an indication of how data augmentation aids in identifying the accurate number of profiles.

To further assess the effect of data augmentation, we also computed the average Precision, Recall, and F1 score across the five scenarios, varying the cosine similarity threshold from 0.8 to 1.

In particular, Fig 3 shows that the overall performance, notably the sensitivity for signature profile detection, improves with the size of data augmentation. This confirms that employing data augmentation as a strategy allows for detecting signature profiles with very high accuracy. Despite the method proving to be fairly accurate even at lower levels of data augmentation, the results clearly illustrate how the use of data augmentation can serve as an effective technique to enhance extraction performance further.

**Figure 3:**
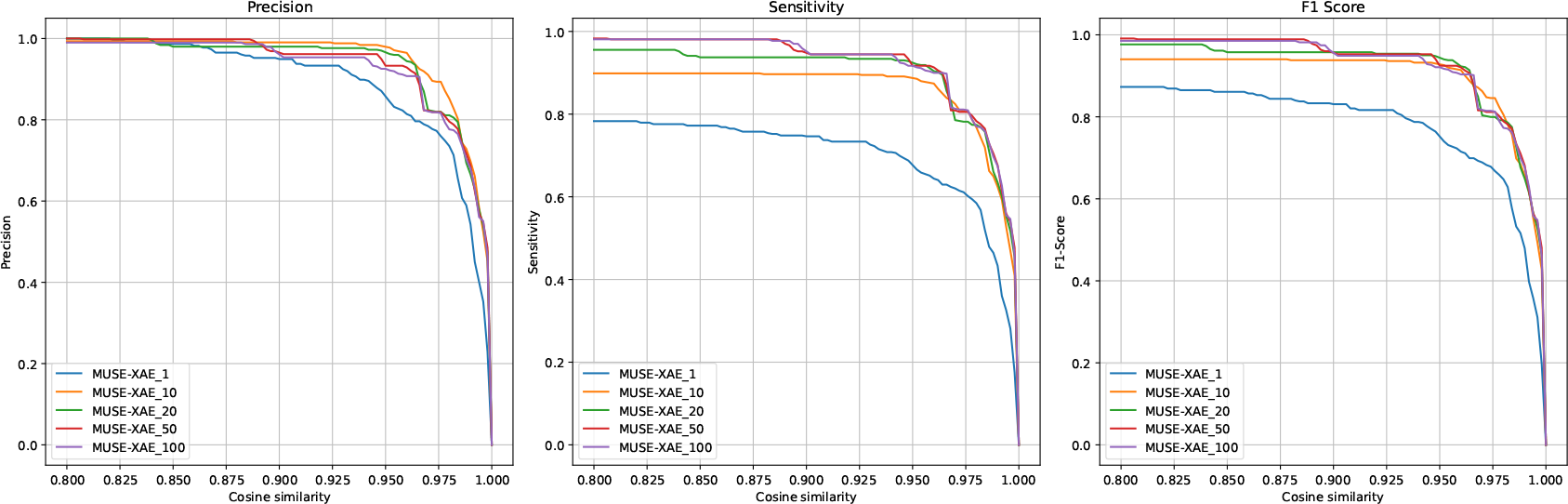
Average Precision, Sensitivity and F1-Score across five scenarios for MUSE-XAE at different level of data augmentation and varying the cosine similarity threshold from 0.8 to 1.

**Figure 4:**
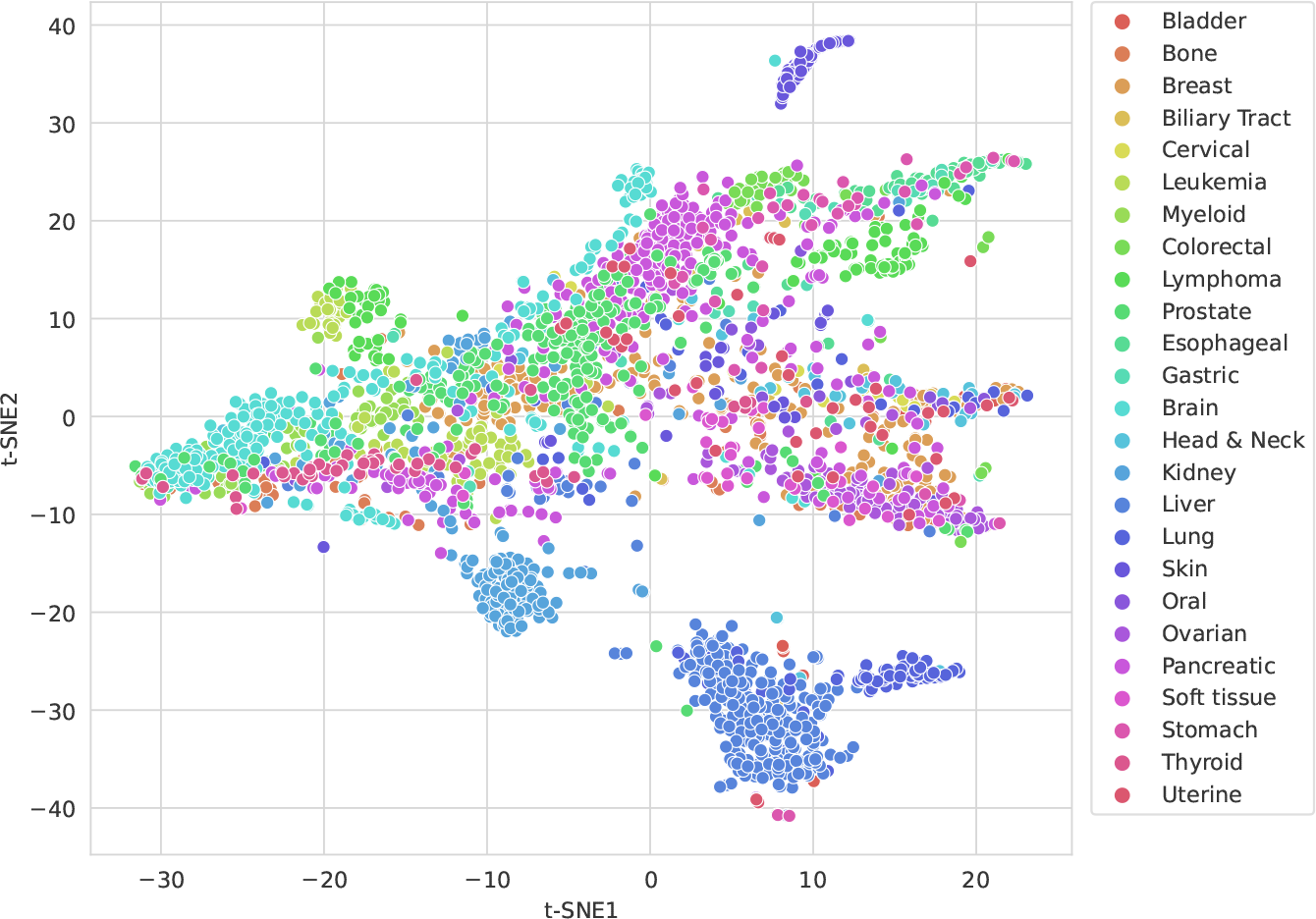
t-SNE representation of the latent representation of the PCAWG datasets, post hoc coloured by primary tumour types

### Performance Comparison

To better establish the performance of our novel method, we compared MUSE-XAE (that uses 100 data augmentation) with 10 existing tools for De-Novo extraction of mutational signatures. The signatures extracted by the other 10 tools across the five scenarios were downloaded from ftp://alexandrovlab-ftp.ucsd.edu/pub/publications/Islam_et_al_SigProfilerExtractor/. To analyze the accuracy of the extracted signatures profiles, we computed Precision, Sensitivity, and F1-score for each scenario, varying the cosine similarity threshold between the extracted and the real profile from 0.8 to 1 for each method.

Supplementary Fig. S1 presents Precision, Sensitivity and Recall of the top 10 performing methods (based on the F1-score), averaged across the five scenarios and Supplementary Fig. S2 the F1-score varying with the cosine similarity threshold for each scenario separately.

Supplementary Fig. S2 shows how each scenario has a different best-performing method, which is generally contested among MUSE-XAE, SigProfilerExtractor, SigProfilerPCAWG, and SigneR. However, globally, Fig. S1 and Table 1 reveal that MUSE-XAE in this five synthetic scenario is the best performing method in all metrics, followed by SigProfilerExtractor and SigProfilerPCAWG.

**Table 1:**
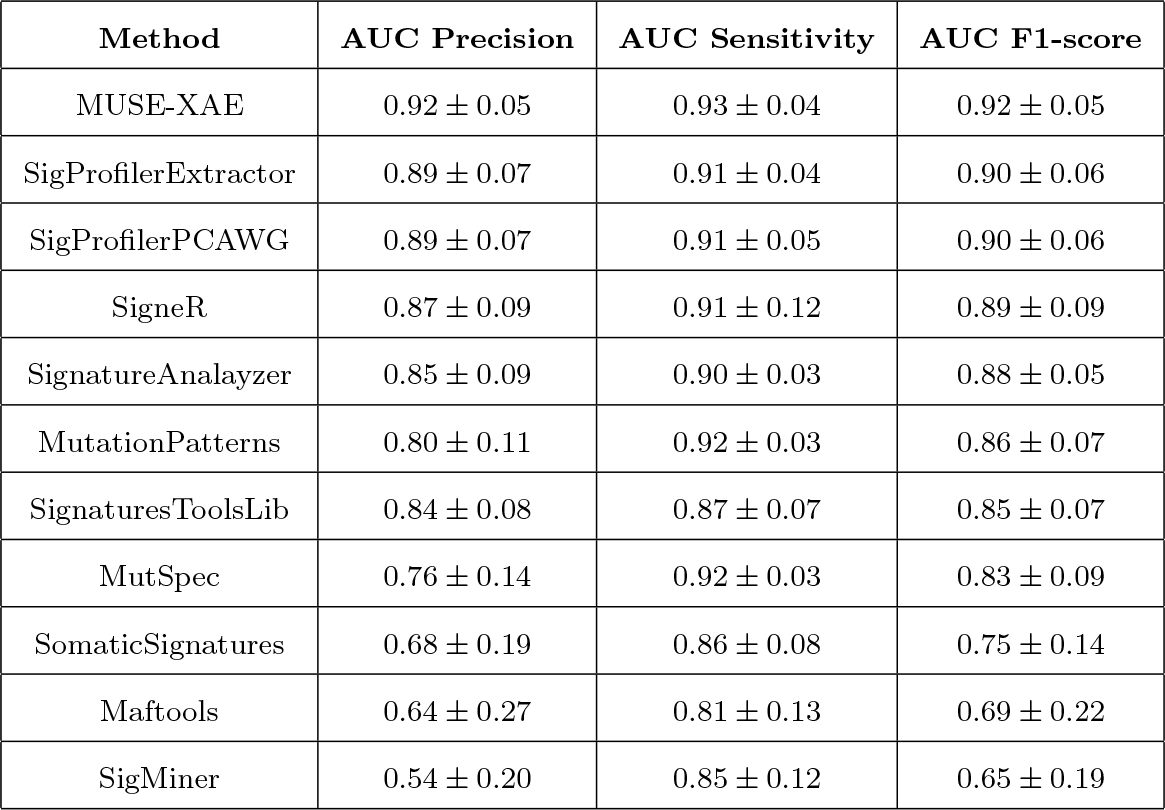
*AUC*_*norm*_ for sensitivity, precision and F1 score curves for each method averaged across the five synthetic scenarios. Methods are ordered based on AUC F1-score.

| Method | AUC Precision | AUC Sensitivity | AUC F1-score |
| --- | --- | --- | --- |
| MUSE-XAE | $0.92 \pm 0.05$ | $0.93 \pm 0.04$ | $0.92 \pm 0.05$ |
| SigProfilerExtractor | $0.89 \pm 0.07$ | $0.91 \pm 0.04$ | $0.90 \pm 0.06$ |
| SigProfilerPCAWG | $0.89 \pm 0.07$ | $0.91 \pm 0.05$ | $0.90 \pm 0.06$ |
| SigneR | $0.87 \pm 0.09$ | $0.91 \pm 0.12$ | $0.89 \pm 0.09$ |
| SignatureAnalyzer | $0.85 \pm 0.09$ | $0.90 \pm 0.03$ | $0.88 \pm 0.05$ |
| MutationPatterns | $0.80 \pm 0.11$ | $0.92 \pm 0.03$ | $0.86 \pm 0.07$ |
| SignaturesToolsLib | $0.84 \pm 0.08$ | $0.87 \pm 0.07$ | $0.85 \pm 0.07$ |
| MutSpec | $0.76 \pm 0.14$ | $0.92 \pm 0.03$ | $0.83 \pm 0.09$ |
| SomaticSignatures | $0.68 \pm 0.19$ | $0.86 \pm 0.08$ | $0.75 \pm 0.14$ |
| Maftools | $0.64 \pm 0.27$ | $0.81 \pm 0.13$ | $0.69 \pm 0.22$ |
| SigMiner | $0.54 \pm 0.20$ | $0.85 \pm 0.12$ | $0.65 \pm 0.19$ |

### PCAWG analysis

We applied MUSE-XAE for the extraction of mutational signatures from the PCAWG dataset, comprising 2780 samples of 37 different cancer types. In particular, we used MUSE-XAE with the data augmentation strategy (100 times the original dataset size for 50 iterations) to find stable consensus signatures. MUSE-XAE found 22 mutational signature profiles, which are presented in Supplementary Fig. S3.

By comparing the 22 profiles identified by MUSE-XAE with the 21 found by SigProfilerExtractor, we discovered considerable overlap. Specifically, by solving the linear assignment problem, i.e matching MUSE-XAE signatures with ones of SigProfilerExtractor, it resulted that the two methods extracted 21 highly similar profiles, showing a mean cosine similarity of 0.97, with a minimum of 0.89. This suggests a significant agreement between the two methods in identifying signature profiles. In Supplementary Table 2. we present the pairwise cosine similarity between the mutational signatures extracted by the two methods.

**Table 2:** Classification performance of MUSE XAE and SigProfilerExtractor using MCC, Kappa, and Balance Accuracy for 27 primary tumour type classification.

| Model | Matthews correlation | Cohen kappa score | Balance Accuracy |
| --- | --- | --- | --- |
| MUSE-XAE | $0.73 \pm 0.02$ | $0.73 \pm 0.02$ | $0.56 \pm 0.03$ |
| SigProfilerExtractor | $0.69 \pm 0.01$ | $0.69 \pm 0.01$ | $0.54 \pm 0.01$ |

However, we discovered differences by matching MUSE-XAE and SigProfilerExtractor signatures with those of the COSMIC database.

In particular, out of the 22 signatures identified by MUSEXAE, 18 exhibit a cosine similarity greater than 0.8 when compared with COSMIC signatures. Similarly, 17 out of the 21 signatures identified by SigProfilerExtractor also demonstrate a cosine similarity greater than 0.8 with COSMIC signatures. Differences and similarities among MUSE-XAE signatures that matched with COSMIC ones are highlighted in Supplementary Figure 4.

From the data presented in the figure, it can be observed that 13 of the COSMIC-matched signatures are present both in MUSE-XAE and SigProfilerExtractor. At the same time, 5 are unique to MUSE-XAE, such as SBS92 associated with tobacco smoking, SBS38 associated with an indirect effect of ultraviolet exposures, SBS26 associated with defective DNA mismatch repair, SBS 60 linked to a sequencing artefact and SBS23 with unknown etiology, and 4 are unique to SigProfilerExtractor, such as SBS45 and SBS54 both linked to sequencing artefact, and SBS16 and SBS19 with unknown etiology.

Among the four MUSE-XAE profiles with a cosine similarity less than 0.8, two have similarities of 0.78 and 0.79 with COSMIC SBS29 and SBS34, respectively. This might suggest an incomplete extraction of these two signatures maybe due to low contribution on mutational profile or due to a concurrence with other signatures. The other two profiles (MUSE-XAE SBS6 and SBS9) fall under the 0.75 of cosine similarity which is the cosine similarity among two 96-mutational channel random profiles.

Among them, we discovered that MUSE-XAE SBS9 has a cosine similarity of 0.86 with the SBS103 of the Signal database [13], a signature with an unknown etiology found both in the ICGC and GEL cohorts [29].

For MUSE-XAE SBS6 we found no matching in any other databases. This signature displays a profile with mutations across many of the 96 mutational channels, with various peaks primarily of *C > A, T > A*, and *T > G* mutational classes. A deeper investigation should be conducted in the future to asses if it is a new signature never reported or if it may represent a noise fitting.

Finally, to thoroughly evaluate the performance of MUSEXAE, we examined the exposure of mutational signatures, i.e the latent representation *z* of tumor samples. While acknowledging that tumors of the same type may demonstrate a degree of heterogeneity, we hypothesize that these exposures, which indicate the number of mutations caused by a signature within a particular sample, could serve as a key discriminant between different tumor types and subtypes.

In Fig.5 we present the t-distributed stochastic neighbor embedding (t-SNE) of the latent representation coloured by 25 primary tumour types, while in Supplementary Figure S5 the t-SNE representation coloured by the 37 tumour subtypes.

**Figure 5:**
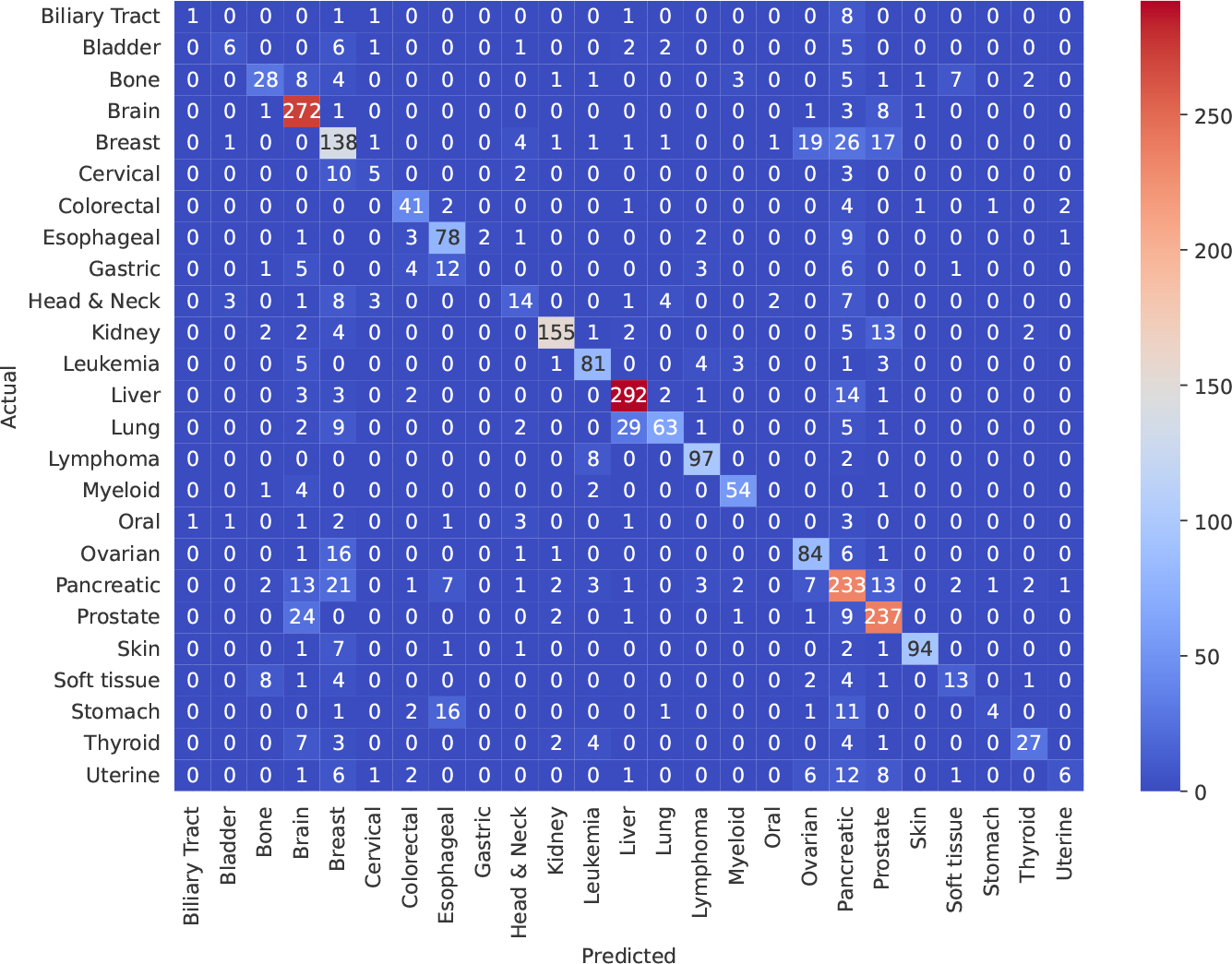
MUSE-XAE Confusion matrix for primary tumour types classification from the latent representation

The t-SNE of exposures displays a clear grouping pattern, which provides compelling evidence supporting this hypothesis, indicating a coherent relationship between signatures exposures and tumor types.

To quantitatively assess this hypothesis, we implemented a Random Forest Classifier that utilizes signature exposures as input to classify both primary tumour types and tumour subtypes. This classifier was applied both to MUSE-XAE and SigProfilerExtractor exposures using a 5-fold cross-validation approach. Confusion Matrix and classification performance metrics for primary tumour types are presented ine Fig.5 and Table 2, while for tumour subtypes in Supplementary Fig S6 and Table S3.

In both classification tasks, MUSE-XAE outperformed SigProfilerExtractor across all metrics, suggesting that the exposures and corresponding signature profiles generated by MUSE-XAE are more discriminative and capable of accurately identifying tumor types.

## 4. Discussion

This paper introduced MUSE-XAE, a novel method for extracting mutational signatures based on autoencoders. Through data augmentation, we implemented a method that demonstrates high accuracy in the De Novo extraction of mutational signatures, proven through a sensitivity analysis and a detailed comparison with 10 other available tools. In particular, MUSE-XAE resulted in the best-performing and robust method in different realistic synthetic scenarios.

Additionally, we analyzed the PCAWG dataset by conducting a de novo extraction of mutational signatures. MUSE-XAE identified 22 mutational signature profiles, exhibiting a high degree of alignment with the profiles extracted by the state-ofthe-art method, SigProfilerExtractor. However, we highlighted discrepancies both in the quantity - MUSE-XAE extracts one additional profile - and in the match with the signatures of the COSMIC database. The fact that 21 profiles of MUSE-XAE and SigProfilerExtractor align with a high degree of similarity yet concurrently match with different COSMIC reference signatures is most likely due to the inherent similarity among several signatures within the COSMIC database, as indicated by various authors. Therefore, two highly similar profiles could still correspond with two different reference signatures. This issue remains an open problem and necessitates further investigations.

However, the profiles extracted by MUSE-XAE appear to be very informative. Indeed, the classification performance based on the exposures shows an MCC (Matthews Correlation Coefficient) of 0.73 and 0.72 in predicting primary types and tumour subtypes, respectively. This achievement confirms that the signatures are discriminative for recognizing different primary cancer types and subtypes with high accuracy.

Finally, MUSE-XAE opens up new possibilities for the development of interpretable neural network-based models for mutational signature extraction, which can leverage the increasing amount of available data and their scalability for large datasets. Our architecture, given its extreme flexibility, can serve as a foundation for building more sophisticated models that could integrate the profile of somatic mutations with other clinical and genomic information, potentially improving and refining the extraction of mutational signatures.

## 6 Author contributions statement

C.P, P.F and T.S. conceived the experiment(s), C.P conducted the experiment(s), C.P, C.R and G.B analysed the results. All the authors wrote and reviewed the manuscript.

## 7 Acknowledgments

We would like to thank Professor Anders Krogh for his invaluable insights and discussions that have significantly improved the quality of the paper

