## Supplementary materials for "MUSE-XAE: MUtational Signature Extraction with eXplainable AutoEncoder enhances tumour type classification"

Corrado Pancotti, Cesare Rollo, Giovanni Birolo, Tiziana Sanavia, Piero Fariselli

October 23, 2023

Table 1: $AUC_{norm}$ for F1 score curves of each method averaged across the five synthetic scenarios

| **Method** | **AUC F1** |
| --- | --- |
| MUSE-XAE | $0.92 \pm 0.05$ |
| SigProfilerPCAWG | $0.90 \pm 0.06$ |
| SigProfilerExtractor | $0.90 \pm 0.06$ |
| SigneR | $0.89 \pm 0.09$ |
| SignatureAnalayzer | $0.88 \pm 0.05$ |
| MutationPatterns | $0.86 \pm 0.07$ |
| SignaturesToolsLib | $0.85 \pm 0.07$ |
| MutSpec | $0.83 \pm 0.09$ |
| SomaticSignatures | $0.75 \pm 0.14$ |
| Maftools | $0.69 \pm 0.22$ |
| SigMiner | $0.65 \pm 0.19$ |

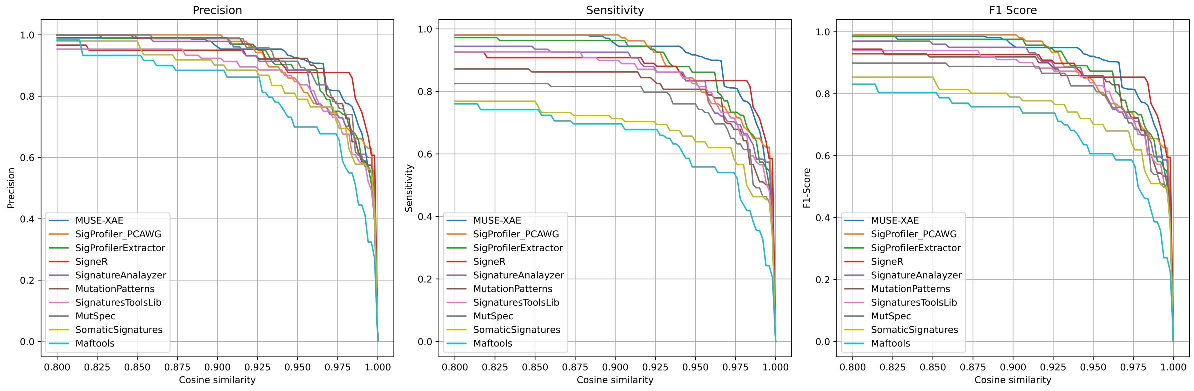


Figure 1: Comparison performance between the top 10 performing methods. On the x-axis cosine similarity thresholds, while on y axes mean Precision, Sensitivity and F1-Score across the five synthetic scenarios. Methods are ordered by the F1 score AUC.

| MUSE-XAE | SigProfilerExtractor | Similarity |
|---|---|---|
| SBS 1 | SBS96I | 0.91 |
| SBS 2 | SBS96R | 0.98 |
| SBS 4 | SBS96E | 0.97 |
| SBS 5 | SBS96H | 1.00 |
| SBS 6 | SBS96N | 0.99 |
| SBS 7 | SBS96O | 0.99 |
| SBS 8 | SBS96G | 0.98 |
| SBS 9 | SBS96Q | 0.93 |
| SBS 10 | SBS96L | 0.99 |
| SBS 11 | SBS96S | 0.94 |
| SBS 12 | SBS96B | 1.00 |
| SBS 13 | SBS96P | 0.99 |
| SBS 14 | SBS96K | 0.98 |
| SBS 15 | SBS96F | 0.95 |
| SBS 16 | SBS96U | 1.00 |
| SBS 17 | SBS96M | 0.94 |
| SBS 18 | SBS96C | 0.99 |
| SBS 19 | SBS96A | 0.99 |
| SBS 20 | SBS96D | 1.00 |
| SBS 21 | SBS96J | 0.99 |
| SBS 22 | SBS96T | 0.89 |

Table 2: Cosine Similarity between the 21 most similar MUSE-XAE and SigProfilerExtractor mutational signatures extracted from the PCAWG dataset

| Model | Matthews correlation | Cohen kappa score | Balance Accuracy |
|---|---|---|---|
| MUSE-XAE | $0.72 \pm 0.01$ | $0.72 \pm 0.01$ | $0.55 \pm 0.02$ |
| SigProfilerExtractor | $0.67 \pm 0.01$ | $0.67 \pm 0.01$ | $0.49 \pm 0.01$ |

Table 3: Classification performance of MUSE XAE and SigProfilerExtractor using MCC, Kappa, and Balance Accuracy for 37 tumour subtypes classification.

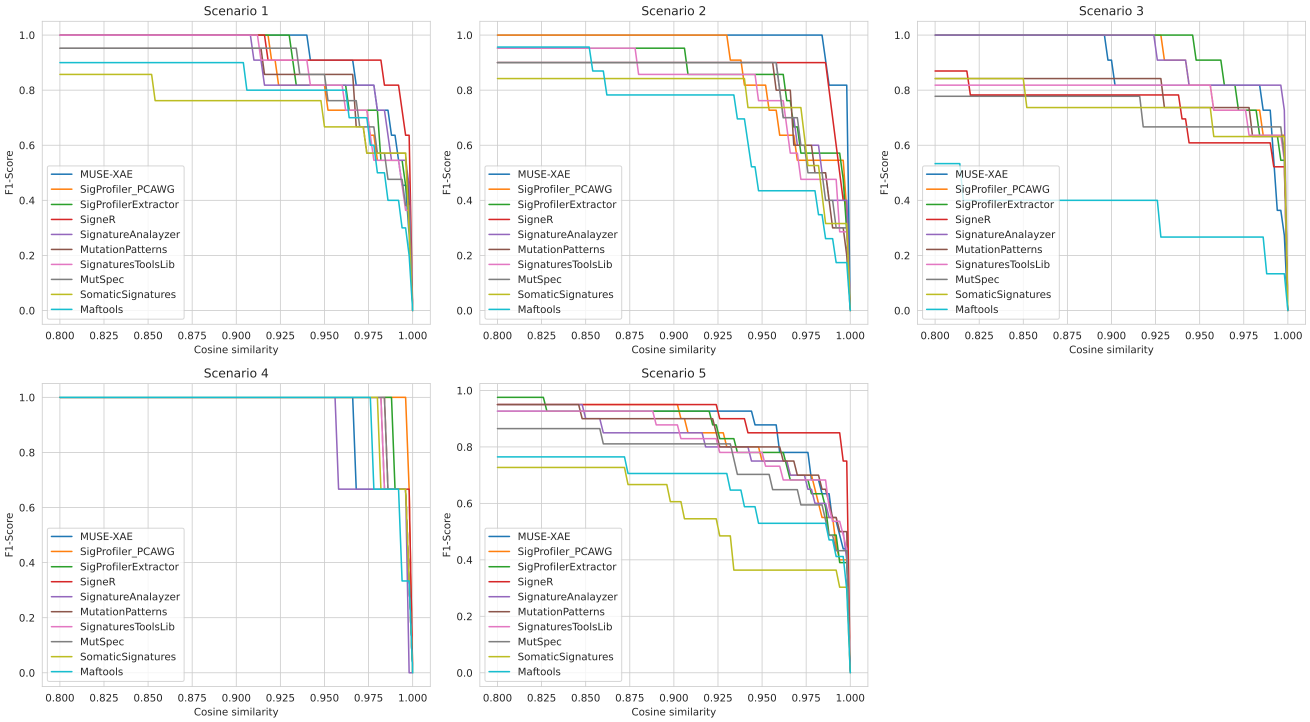


Figure 2: Comparison performance between the top 10 performing methods. On the x-axis cosine similarity thresholds, while on y F1-Score for each synthetic scenario. Methods are ordered by the F1 score AUC.

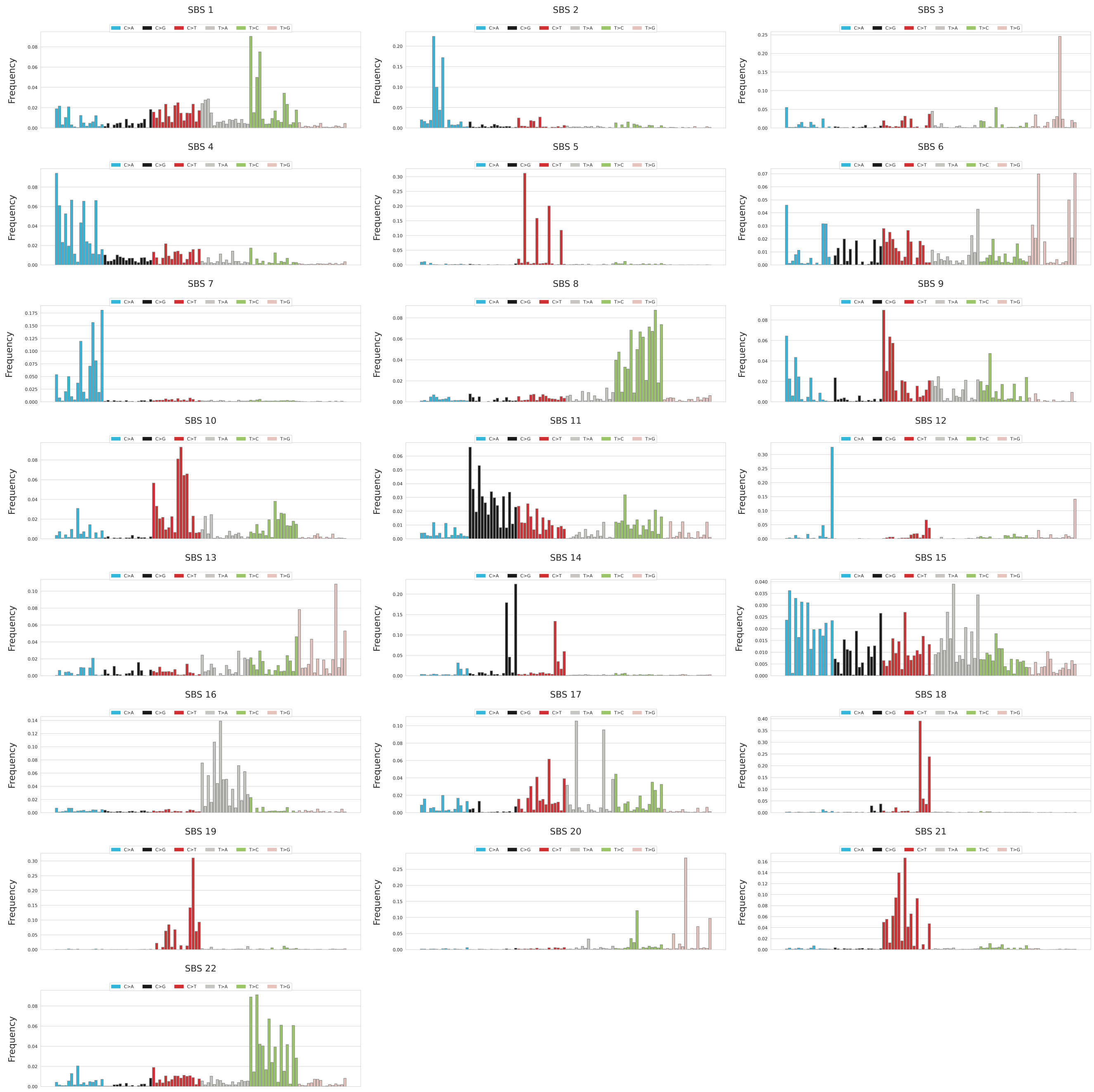


Figure 3: 22 mutational signatures extracted from PCAWG dataset

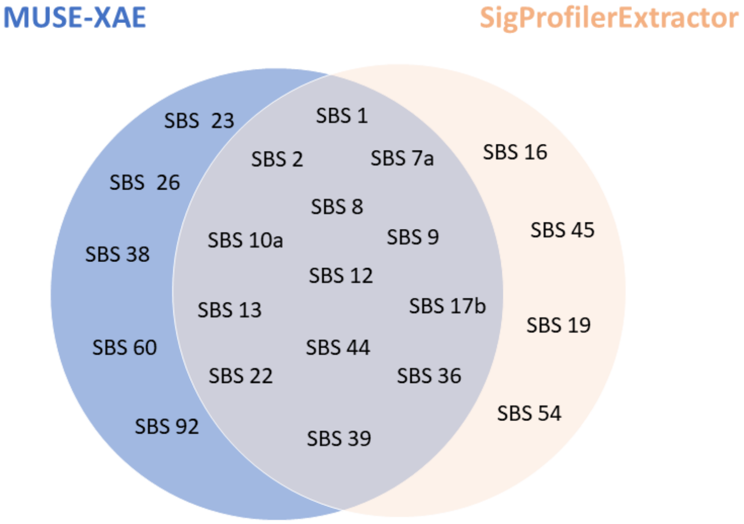


Figure 4: MUSE-XAE and SigProfilerExtractor COSMIC matched signatures on the PCAWG dataset

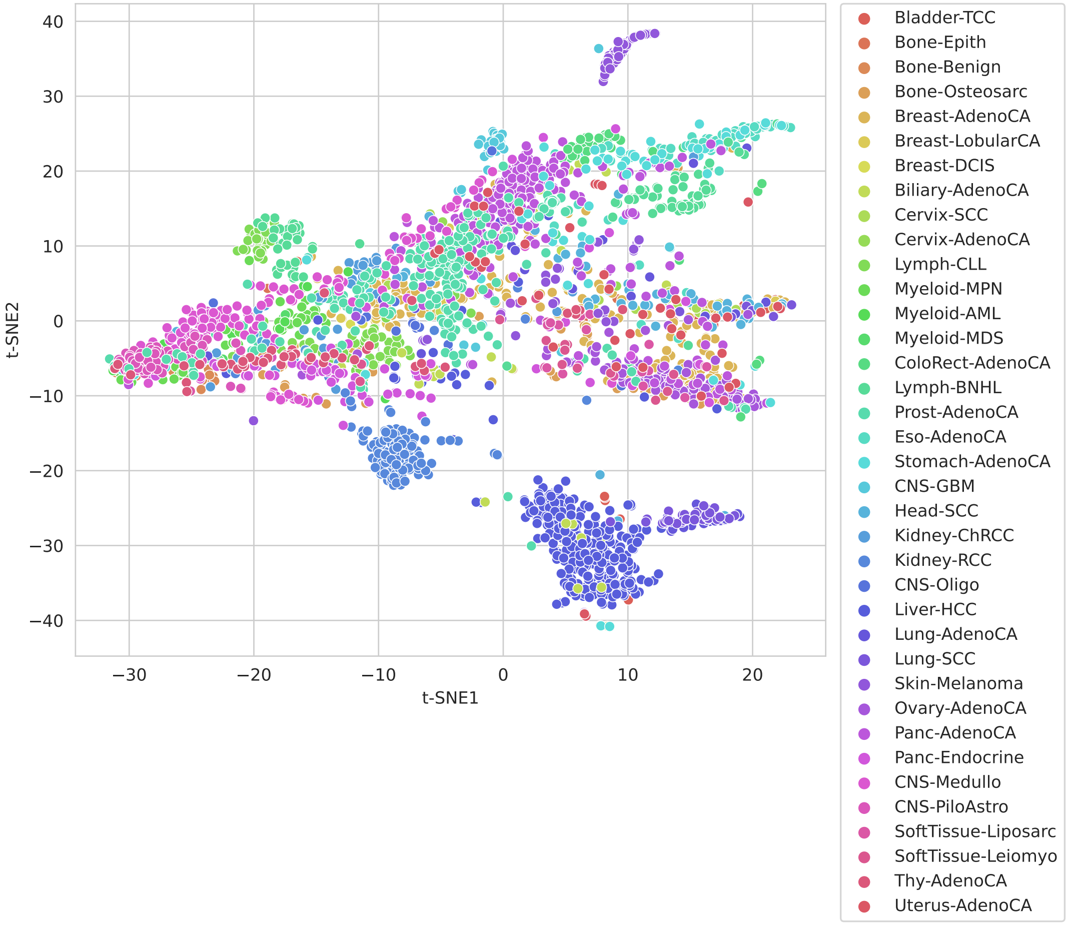


Figure 5: t-SNE representation of the latent representation of the PCAWG datasets, post hoc coloured by tumour subtypes

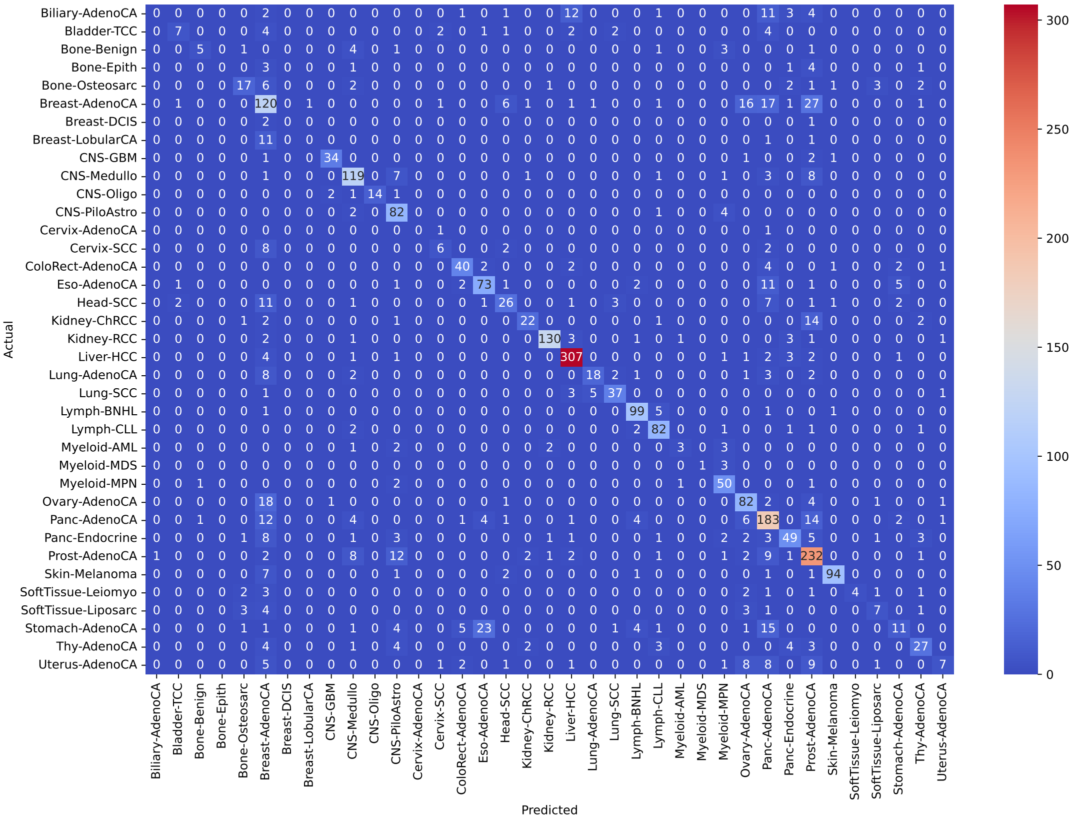


Figure 6: MUSE-XAE Confusion matrix for tumour subtypes classification
